# P2RX7 signaling drives the differentiation of Th1 cells through metabolic reprogramming for aerobic glycolysis

**DOI:** 10.1101/2022.10.24.513424

**Authors:** Érika Machado de Salles, Paulo Lisboa Raeder, Claudia Angeli, Verônica Feijoli Santiago, Cristiane Naffah de Souza, Theresa Ramalho, Niels Olsen Saraiva Câmara, Giuseppe Palmisano, José Maria Álvarez, Maria Regina D’Império Lima

**Affiliations:** Departamento de Imunologia, Instituto de Ciências Biomédicas, Universidade de São Paulo, São Paulo, Brazil; Departamento de Parasitologia, Instituto de Ciências Biomédicas, Universidade de São Paulo, São Paulo, Brazil

## Abstract

This study provides evidence on the molecular mechanisms by which P2RX7 signaling promotes the differentiation of Th1 cells. *In vivo* analysis was performed in the *Plasmodium chabaudi* model of malaria in view of the great relevance of this infectious disease for human health, as well as the great availability of data concerning Th1/Tfh differentiation. We show that P2RX7 induces T-bet expression and aerobic glycolysis in splenic CD4^+^ T cells that respond to malaria, at a time prior to Th1/Tfh polarization. Cell-intrinsic P2RX7 signaling sustains the glycolytic pathway and causes bioenergetic mitochondrial stress in activated CD4^+^ T cells. We also show *in vitro* the phenotypic similarities of Th1-conditioned CD4^+^ T cells that do not express P2RX7 and those in which the glycolytic pathway is pharmacologically inhibited. In addition, *in vitro* ATP synthase blockade and the consequent inhibition of oxidative phosphorylation, which drives cellular metabolism for aerobic glycolysis, is sufficient to promote rapid CD4^+^ T cell proliferation and polarization to the Th1 profile in the absence of P2RX7. These data demonstrate that P2RX7-mediated metabolic reprograming for aerobic glycolysis is a key event for Th1 differentiation and suggest that ATP synthase inhibition is a downstream effect of P2RX7 signaling that potentiates the Th1 response.

## INTRODUCTION

In the last decades, the relevance of purinergic signaling to T cell function has received increasing attention from the scientific community. Adenosine triphosphate (ATP) has been identified as a key molecule for communication between dendritic cells and T cells (Schenk et al., 2008; Woehrle et al., 2010). ATP released by pannexin-1 hemichannels at immunological synapses activates ATP-gated P2RX cation channels on T cells and promotes the entry of Ca^++^. Environmental ATP released by stressed, damaged or dying cells from surrounding tissue as a danger signal can also increase T cell responses. Ca^++^ influx leads to translocation of the nuclear factor of activated T cells (NFAT) to the cell nucleus and consequent production of IL-2. IL-2 signaling induces T cell proliferation and drives the differentiation of T helper 1 (Th1) cells together with some signals form dendritic cells, such as IL-12, and produce IFNγ that activate macrophages. Th1 differentiation depends on the expression of transcriptional factors T-box expressed in T cells (T-bet) and B-lymphocyte-induced maturation protein 1 (Blimp-1) (Gong and Malek, 2007; Liao et al., 2011). As a regulatory mechanism, activated T cells express CD39 and CD73 ectonucleotidases that hydrolyze extracellular ATP (eATP) and generate adenosine, which exerts its regulatory activity through the engagement of G-coupled P1 receptors (Takenaka et al., 2016).

Among the seven members of the P2RX family, T cells express P2RX1, P2RX4 and P2RX7 (Schenk et al., 2008). The latter receptor is particularly involved in pathological conditions as it is activated only at high concentrations of eATP and induces the formation of large pores in the cell membrane. The massive influx of Ca^++^ through P2RX7-induced membrane pores can potentiate cell activation or lead to cell death under prolonged stimulus (Steinberg and Silverstein, 1987; Steinberg et al., 1990). As an evidence of these effects, P2RX7 is upregulated in CD4^+^ T cells activated during acute murine malaria caused by *Plasmodium chabaudi* and promotes membrane pore formation, intense Ca^++^ influx and IL-2 production, and thus induces a vigorous Th1 response that protects mice from death (Salles et al., 2017). Conversely, P2RX7 deficiency in CD4^+^ T cells during acute *P. chabaudi* infection leads to the predominance of early follicular helper T (Tfh) cells expressing the transcriptional factor B-cell lymphoma 6 (Bcl6). Both Th1 and Tfh subsets contribute to protection against *P. chabaudi* infection, as in the human malaria, evidenced by the protective role of IFNγ and parasite-specific antibodies (Perez-Mazliah and Langhorne, 2015). P2RX7 may also participate in the metabolic changes observed in activated T cells, particularly during intense immune responses and tissue damage. However, information on this subject is scarce, mainly addressing the contribution of P2RX7 to the metabolic fitness of long-lived memory CD8^+^ T cells developed after virus infection (Borges da Silva et al., 2018).

T cell subpopulations have metabolic differences that regulate cell fate and function (Chi, 2012; MacIver et al., 2013). Resting and memory T cells oxidize pyruvate, lipids and amino acids to produce the basal level of ATP that is necessary for cell survival (Van der Windt and Pearce, 2012). Ca^++^ input mechanisms, such as the stockade-calcium intake (SOCE), support T cell proliferation and differentiation to effector functions through metabolic reprogramming (Vaeth et al, 2017). After T cell activation, lipid oxidation is downregulated and aerobic glycolysis increases along with glutamine oxidation in order to supply the biosynthetic precursors required for rapid cell proliferation (Carr et al., 2010). Regarding CD4^+^ T cell subsets, Th1 cells are more proliferative and metabolically active than Tfh cells, which induce antibody class switching and affinity maturation in B cells (Ray et al, 2015). In order to maintain an efficient Th1 response, activation of metabolic sensors Akt and the mammalian target of rapamycin (mTOR) drives CD4^+^ T cell fate through upregulation of aerobic glycolysis (Delgoffe et al., 2009). As an indication that the ATP-P2RX7 axis also modulates T cell metabolism, overexpression of P2RX7 confers an intrinsic ability to upregulate aerobic glycolysis and mitochondrial reactive oxygen species (ROS) in fast-growing tumor cell lines (Amoroso et al., 2012). It has been shown that mitochondrial ROS is critical for T cell activation by inducing NFAT translocation to the cell nucleus and subsequent IL-2 production (Kamiński et al., 2010; Sena et al., 2013).

Here, we investigated how P2RX7 modulates CD4^+^ T cell metabolism during strong Th1 responses. *In vivo* studies were performed in *P. chabaudi*-infected mice in view of the great relevance of malaria for human health, as well as the large amount of information available by flow cytometry and single cell RNAseq analysis concerning Th1/Tfh polarization in this experimental model (Muxel et al., 2011; Salles et al., 2017; Lönnberg et al., 2017; Soon et al., 2020). The single cell RNAseq analysis highlights the metabolic profile as a determining factor for the fate of malaria-responsive CD4^+^ T cells during the Th1/Tfh subset bifurcation, as cells cycling fastest and exhibiting the greatest glycolytic activity are the first to acquire the ability to produce IFNγ (Lönnberg et al., 2017). Confirmation and further details on the effects of P2RX7 on CD4^+^ T cell metabolism were obtained by *in vitro* analysis under intense Th1 polarization. Our results demonstrate that P2RX7-induced aerobic glycolysis is a key event for the development of efficient Th1 responses. P2RX7 signaling in CD4^+^ T cells upregulates Akt and mTOR, as well as glucose uptake, promoting proliferation and differentiation of Th1 cells. Inducing the production of ATP synthase inhibitors appears to be an important molecular mechanism by which P2RX7 signaling in CD4^+^ T cells upregulates aerobic glycolysis and, therefore, Th1 differentiation.

## RESULTS

### P2RX7 induces early upregulation of T-bet transcriptional factor in CD4^+^ T cells that respond to *Plasmodium* infection

Previous studies have shown that Th1/Tfh subsets emerge in parallel by day 7 post-infection (p.i.) with *P. chabaudi* in B6 mice (Salles et al., 2017), but subset bifurcation revealed by gene expression signatures occurs earlier around day 4 p.i. (Lönnberg et al., 2017). Because P2RX7 has a fundamental role in the generation of Th1 cells during experimental malaria (Salles et al., 2017), we sought to determine the first cellular events through which this molecule supports the Th1 response. Initially, the expression of Blimp1, T-bet and Bcl6 transcriptional factors was evaluated in CD4^+^ T cells from B6 and *P2rx7*^-/-^ mice at the acute phase of infection. These cells were stained for FoxP3 to exclude regulatory T cells in the phenotypic analysis. A single population of Foxp3^-^CD4^+^ T cells co-expressed T-bet and Bcl6 at days 5 and 6 p.i., with comparable numbers of these cells per spleen being found in B6 and *P2rx7*^-/-^ mice (Figure 1A). Of note, T-bet expression was lower in Foxp3^-^CD4^+^ T cells from *P2rx7*^-/-^ mice at day 6 p.i., while Bcl6 and Blimp-1 were similarly upregulated in cells of both mouse groups (Figure 1B). As previously reported (Salles et al., 2017), T-bet and Bcl6 expression distinguished Th1 (T-bet^hi^Bcl6^lo^) cells from early Tfh (T-bet^lo^Bcl6^hi^) cells in B6 mice at day 7 p.i., and a reduction in the Th1 subset was observed in *P2rx7*^-/-^ mice (Figure 1C). These findings indicate that P2RX7-induced T-bet upregulation is an early event that precedes Th1/Tfh polarization in murine malaria.

**Figure 1.**
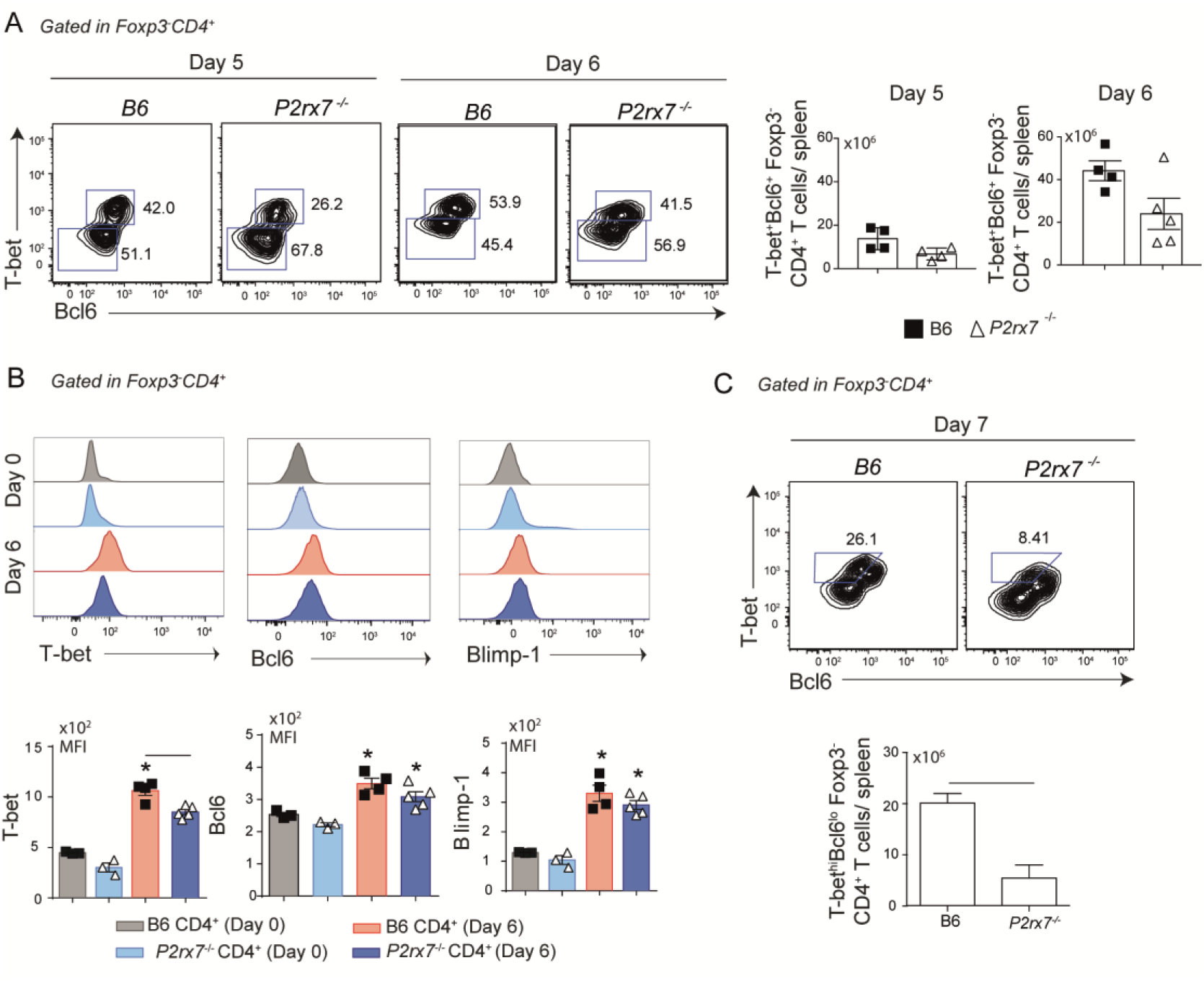
Early effects of P2RX7 on T-bet expression by splenic CD4^+^ T cells that respond to *Plasmodium* infection. B6 and *P2rx7*^-/-^ mice were infected i.p. with 1 × 10^6^ iRBCs. (A) Contours plots show T-bet and Bcl6 expression in Foxp3^-^CD4^+^ T cells at days 5 and 6 of infection. Column bar graphs show T-bet^+^Bcl6^+^Foxp3^-^CD4^+^ T cell numbers per spleen. (B) Histograms show T-bet, Bcl6 and Blimp-1 expression in Foxp3^-^CD4^+^ T cells at day 6 of infection. Column bar graphs show median fluorescence intensities (MFIs) for T-bet, Bcl6 and Blimp-1. (C) Contours plots show T-bet and Bcl6 expression in Foxp3^-^CD4^+^ T cells at day 7 of infection. Column bar graph shows T-bet^hi^Bcl6^lo^Foxp3^-^CD4^+^ T cell numbers per spleen. Data are shown as the mean ± SD (n = 4-5) of one representative experiment out of three. Significant differences are observed for the (-) indicated groups with *p* < 0.05, using Mann Whitney U test.

### P2RX7 supports the Th1 response during experimental malaria by activating the glycolytic pathway

In *P. chabaudi* malaria, CD4^+^ T cells proliferate intensely and are metabolically very active around day 4 p.i. and progressively stop dividing and differentiate into effector cells until day 7 p.i., after which most activated cells die in parallel with control of acute parasitemia (Elias et al., 2005; Muxel et al., 2011; Lönnberg et al., 2017). To investigate whether metabolic reprogramming of CD4^+^ T cells by P2RX7 is a key event for Th1 lineage differentiation, we performed a proteomic analysis on Foxp3^-^ CD11a^+^CD4^+^ T cells from B6 and *P2rx7*^-/-^ mice at day 6 of infection. CD11a interacts with ICAM-1 and has been previously used to identify *Plasmodium*-specific T cells responding to infection (Butler et al., 2011). Either CD11a^+^ cell populations, expressing or not P2RX7, were activated as evidenced by their large size (FSC-A) and positivity for T-bet and Bcl6, and were present in similar percentages among FoxP3^-^ CD4^+^ T cells (Figure 2A). However, B6 CD4^+^ T cells were larger than *P2rx7*^-/-^ CD4^+^ T cells (Figure 2B), which is indicative of increased proliferation and/or differentiation in effector cells in the presence of P2RX7; large cell size is a feature of both proliferating and IFNγ-producing B6 CD4^+^ T cells (Lönnberg et al., 2017).

According to heatmap analysis of label-free quantitative data, 121 of the upregulated proteins were found in *P2rx7*^-/-^ CD4^+^ T cells compared to B6 CD4^+^ T cells and 37 of the upregulated proteins were found in B6 CD4^+^ T cells compared to *P2rx7*^-/-^ CD4^+^ T cells (Figure 2C and Table S1). Higher numbers of upregulated proteins were found in *P2rx7*^-/-^ CD4^+^ T cells for all assessed biological pathways as categorized using the Reactome program, including cell cycle and DNA replication pathways (Figure 2D). Concerning the metabolic pathways, however, proteins involved in the glucose metabolism were selectively increased in B6 CD4^+^ T cells (Figure 2E). Among these proteins, hexokinase 1 (Hk1), triosephosphate isomerase 1 (Tpi1) and fructose-biphosphatealdolase A (Aldoa) play a key role in the glycolytic pathway, as well as in the reverse gluconeogenesis pathway; alpha-enolase (Eno1) catalyzes the conversion of 2-phosphoglycerate to phosphoenolpyruvate required for pyruvate synthesis (Cappello et al., 2017). Expression of glycolytic enzymes was associated with upregulation of isocitrate dehydrogenase subunit α (Idh3a) and ATP synthase subunit β (ATP5f1), which participate in the tricarboxylic acid (TCA) cycle and respiratory electron transport, respectively. In contrast, P2RX7 deficiency led to upregulation of enzymes involved in the synthesis of nucleotides and amino acids, such as phosphoserine aminotransferase (Psat1), D-3-phosphoglycerate dehydrogenase (Phgdh), trifunctional purine biosynthetic protein adenosine-3 (Gart) and inosine-5’ monophosphate dehydrogenase 2 (Impdh2). These proteins were associated with isocitrate dehydrogenase subunit γ (Idh3g) and the F6 subunit of the ATP synthase peripheral stalk (ATP5j). In addition, P2RX7 deficiency promoted the upregulation of several proteins involved in lipid synthesis, such as acyl-CoA thioesterase (Acot7), fatty acid synthase (Fasn) and isopentenyl-diphosphate delta-isomerase 1 (Idi1).

**Figure 2.**
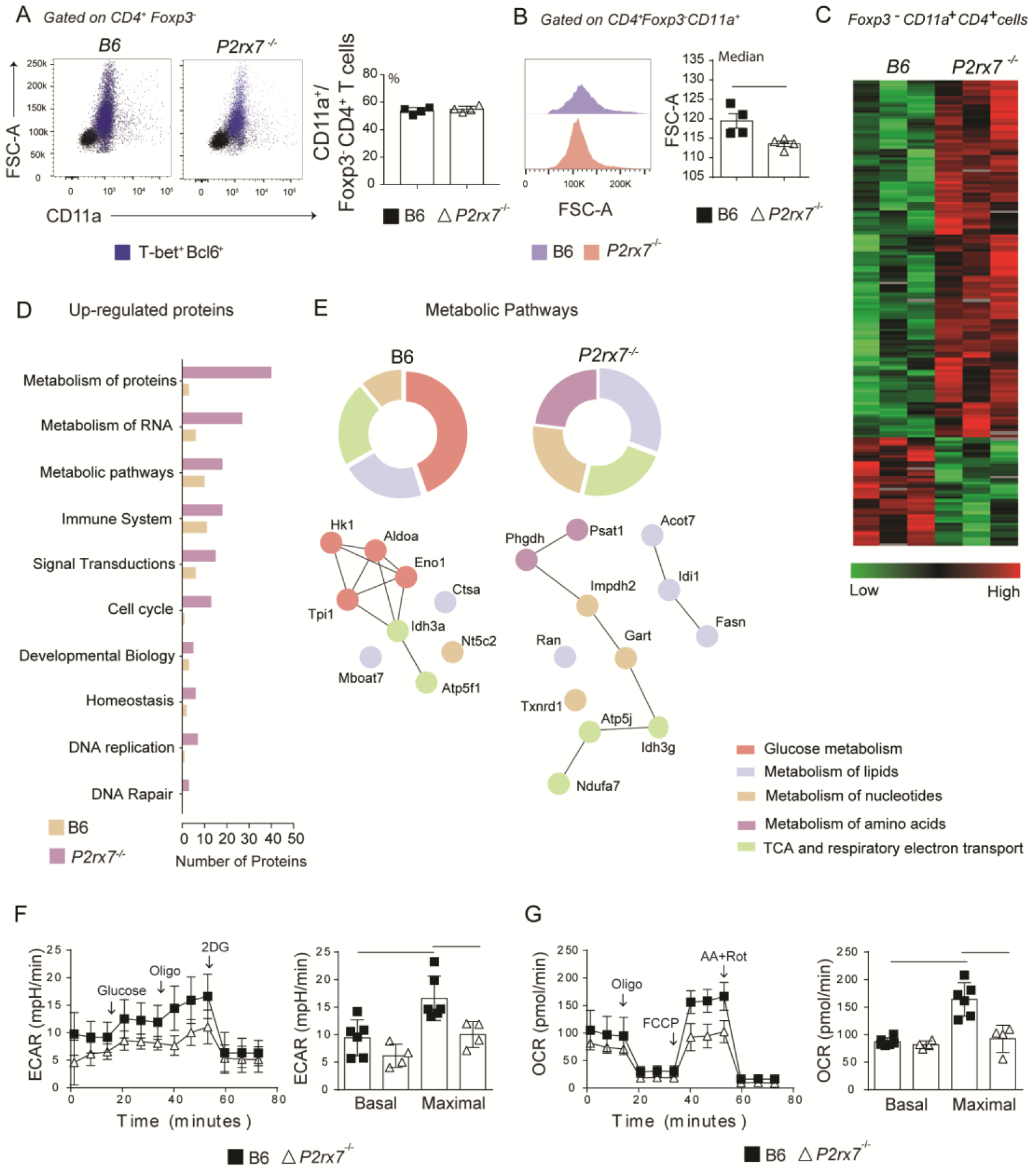
Metabolic analysis of splenic CD4^+^ T cells from B6 and *P2rx7*^-/-^ mice at early *Plasmodium* infection. Foxp3.GFP and Foxp3.GFP^+^*P2rx7*^-/-^ mice were infected i.p. with 1 × 10^6^ iRBCs and analyzed 6 days later. (A) Dot plots show forward side scatter area (FSC-A) and CD11a expression in Foxp3^-^CD4^+^ T cells. Blue dots identify T-bet^+^Bcl6^+^ cells. Column bar graph shows CD11a^+^ cell frequencies in the Foxp3^-^CD4^+^ T cell population. (B) Histograms show FSC-A of CD11a^+^Foxp3^-^CD4^+^ T cells. Column bar graphs show FSC-A medians (C) The heatmap of label-free quantitative data shows upregulated proteins in red and downregulated proteins in green. (D) Functional groups of biological pathways defined by Reactome Database show upregulated proteins. (E) Enrichment of metabolic pathways defined by Reactome Database is shown. The upregulated proteins are provided in the interaction network and their first nodes. (F and G) Real time changes in extracellular consumption acidification rate (ECAR) and oxygen consumption rate (OCR) were measured using a Seahorse XFe96 bioanalyzer. (F) The first arrow indicates the injection of glucose to increase the glycolytic pathway, followed by oligomycin to enhance the glucose consumption (second arrow) and 2-deox-D-glucose (2DG) to inhibit glycolysis (third arrow). (G) The first arrow indicates the injection of oligomycin that promotes a metabolic stress, followed by a mitochondrial decoupler carbonyl cyanide p-trifluoro-methoxyphenyl hydrazone (FCCP) to determine the maximal respiratory capacity (second arrow) and antimycin A/rotenone to block the electron transport chain (third arrow). Data are shown as the mean ± SD (n = 4) of one representative experiment out of three. Significant differences were observed for the (-) indicated groups with p < 0.05, using the Mann Whitney U.

To assess the impacts of P2RX7 expression in the glycolytic and oxidative capacity of CD4^+^ T cells at day 6 p.i., the extracellular consumption acidification rate (ECAR) and oxygen consumption rate (OCR) were evaluated. No difference was observed in basal rates, but maximal ECAR and OCR were reduced in *P2rx7*^-/-^ CD4^+^ T cells compared to B6 CD4^+^ T cells (Figures 2F and 2G). It is noteworthy that inhibition of oxidative phosphorylation by blocking ATP synthase with oligomycin upregulates ECAR in B6 and *P2rx7*^-/-^ CD4^+^ T cells, although the former population showed a greater increase.

According to our proteomics and seahorse data it is possible to suggest that P2RX7 promotes a more energetic state of anabolic profile in the cell by inducing block synthesis pathways. Glycolysis appears to support the availability of substrates for this metabolic state and therefore allow rapid CD4^+^ T cell proliferation and differentiation to the Th1 profile. Otherwise, in the absence of P2RX7, most CD4^+^ T cells that respond to *Plasmodium* infection show an early Tfh profile and maintain a slow but sustained proliferative activity, giving rise to fully differentiated (PD1^hi^CXCR5^+^) Tfh cells around day 14 p.i. (Salles et al., 2017; Soon et al., 2020).

### Cell-intrinsic P2RX7 expression promotes mTOR-coordinated metabolic pathways in CD4^+^ T cells that respond to *Plasmodium* infection

To verify whether P2RX7 expression in CD4^+^ T cells is sufficient to induce the metabolic shift required for Th1 differentiation, splenic CD4^+^ T cells were sorted from naïve B6 (CD45.1^+^) and *P2rx7*^-/-^ (CD45.1^+^) donors and co-transferred to *Cd4*^-/-^ recipients that were infected with *Plasmodium* (Figure 3A). Using this experimental approach, B6 and *P2rx7*^-/-^ CD4^+^ T cells sharing the same environment could be compared. Similar numbers of co-transferred CD45.1^+^CD4^+^ and CD45.2^+^CD4^+^ T cells per spleen were found on day 6 p.i., as well as comparable percentages of CD11a^+^ cells in these populations (Figures S1A and S1B). Remarkably, P2RX7 expression in CD4^+^ T cells increased cell size and proliferation, as revealed by FSC-A and Ki67 staining (Figures 3B and 3C). In addition, most effector CD4^+^ T (CD44^+^CD62L^-^ and CD44^+^CD39^+^CD11a^+^) cells expressed P2RX7 in infected mice (Figures S1C and S1D); CD39 is highly expressed in Th1 cells (Salles et al., 2017). Evidencing the crucial role of P2RX7 for Th1 differentiation, IFNγ-producing CD44^+^CD11a^+^CD4^+^ T cells were overrepresented in the population that expressed this receptor (Figure 3D). Regarding the metabolic profile, P2RX7 expression in CD4^+^ T cells resulted in increased phosphorylation of mTOR and Akt (Figure 3E), and upregulation of glucose uptake as seen by 2-NBDG (2-deoxy-2-[(7-nitro-2,1,3-benzoxadiazol-4-yl)amino]-D-glucose) staining (Figure 3F). Of note, the metabolic change induced in activated CD4^+^ T cells by P2RX7 signaling was associated with increased ROS production as revealed by DCFH (2’,7’-dichlorofluorescein) staining (Figure 3G). In addition, a higher proportion of B6 CD4^+^ T cells showing increased mitochondrial mass also presented mitochondrial depolarization in response to *Plasmodium* infection (Figure 3H). This was evidenced by staining CD11a^+^CD4^+^ T cells with the mitochondrial membrane potential (MMP) indicator TMRE (tetramethylrhodamine ethyl ester) and MTG (MitoTracker Green).

**Figure 3.**
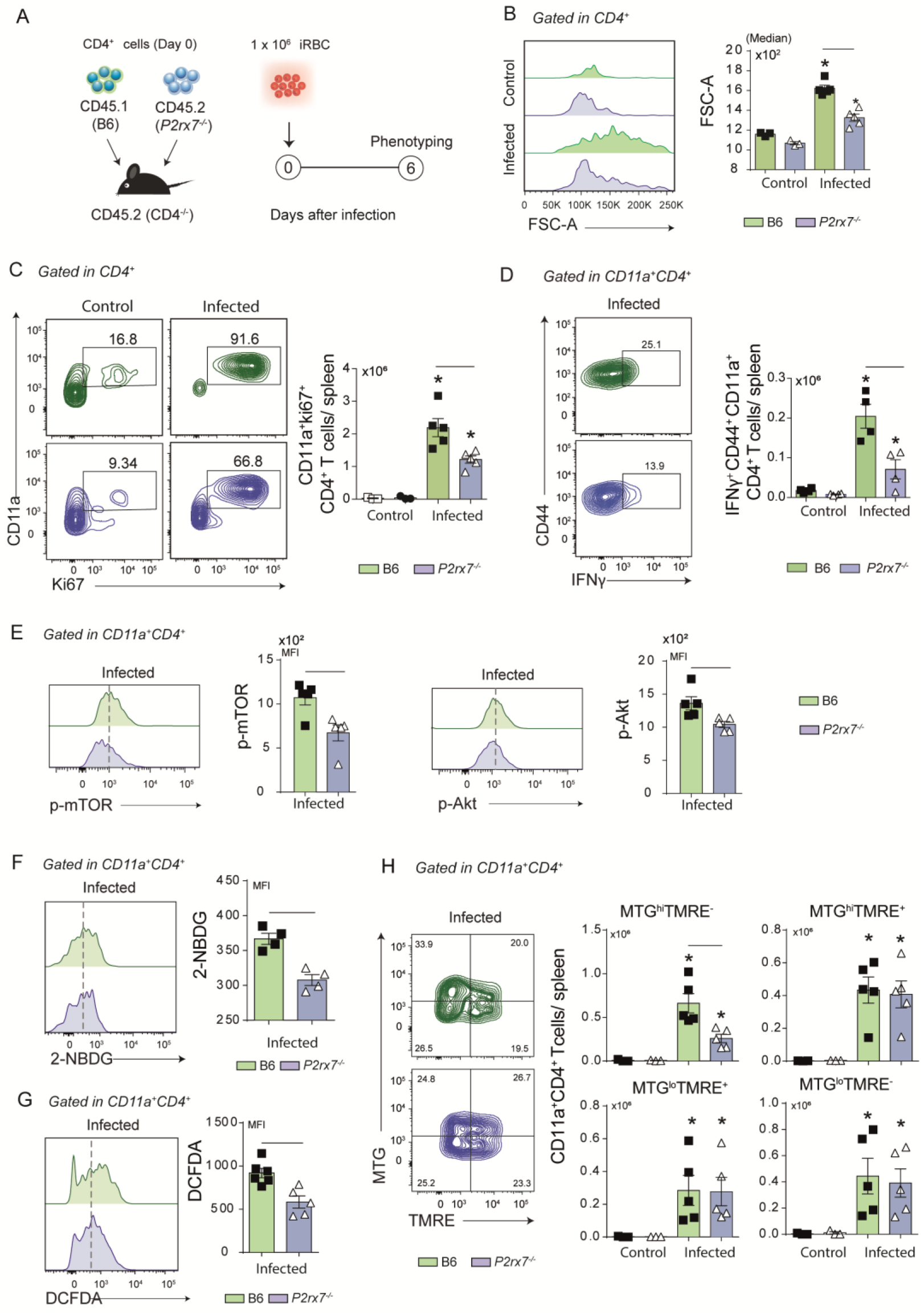
Phenotypical analysis of metabolic markers in co-transferred B6 and *P2rx7*^-/-^ CD4^+^ T cells at early *Plasmodium* infection. Splenic naïve CD4^+^ T cells from B6 (CD45.1^+^) and *P2rx7*^-/-^ (CD45.2^+^) mice were co-transferred into *Cd4*^-/-^ mice that were infected with 1 × 10^6^ iRBCs. Splenic CD45.1^+^ and CD45.2^+^ CD4^+^ T cells were analyzed at day 6 of infection. Non-infected mice were used as controls. (A) A schematic illustration of the experimental protocol is shown. (B) Histogram shows forward side scatter area (FSC-A) of CD4^+^ T cells and the medians are shown in the column bar graph. (C) Contour plots show CD11a *versus* Ki67 expression in CD4^+^ T cells. Column bar graph shows CD11a^+^Ki67^+^CD4^+^ T cell numbers per spleen. (D) Contour plots show CD44 *versus* intracellular IFNγ expression in CD11a^+^CD4^+^ T cells. Column bar graph shows IFNγ^+^CD44^+^CD11a^+^CD4^+^ T cell numbers per spleen. (E) Histograms show p-mTOR or p-Akt expression in CD4^+^ T cells. Column bar graphs show mTOR and Akt MFIs. (F) Histogram shows 2-NBDG staining in CD11^+^CD4^+^ T cells. Column bar graph shows 2-NBDG—MFIs. (G) Histogram shows DCFDA (2’,7’-dichlorofluorescin) staining in CD11^+^CD4^+^ T cells. Column bar graph shows DCFDA MFIs. (H) Contour plots show MTG *versus* TMRE staining in CD11a^+^CD4^+^ T cells. Column bar graphs show MTG^hi^TMRE^+^, MTG^hi^TMRE^-^, MTG^lo^TMRE^+^ and MTG^lo^TMRE^-^ CD11a^+^CD4^+^ T cell numbers per spleen. Data are shown as the mean ± SD (n = 4-5) of one representative experiment out of three. Significant differences were observed for the (-) indicated groups and (*) between infected and control mice with p < 0.05, using Mann Whitney U test.

We concluded that cell-intrinsic P2RX7 expression is sufficient to sustain glucose metabolism, as evidenced by mTOR and Akt phosphorylation and glucose uptake upregulation, and induce Th1 differentiation. Increased ROS production and loss of MMP are signals of bioenergetic mitochondrial stress also induced by P2RX7 expression in CD4^+^ T cells that respond to *Plasmodium* infection.

### ATP synthase inhibition restores Th1 differentiation *in vitro* in the absence of P2RX7

To assess whether P2RX7 is important for Th1 differentiation *in vitro*, B6 and *P2rx7* ^-/-^ CD4^+^ T cells were stimulated with anti-CD3 and anti-CD28 antibodies and polarized to the Th1 profile in conditioning medium containing IL-12, IL-2 and anti-IL-4 antibody; stimulated but unpolarized Th0 cells were used as controls. Under Th1-polarizing condition, only B6 CD4^+^ T cells showed T-bet and Blimp-1 upregulation (Figure 4A), and these cells produced more IFNγ than *P2rx7*^-/-^ CD4^+^ T cells (Figure 4B). P2RX7 was also required for CD4^+^ T cell proliferation in this condition, as evidenced by both FSC-A (Figure 4C) and CFSE staining (Figure S2A). Furthermore, the generation of CD44^+^CD4^+^ T cells coexpressing CD25 (IL-2R) (Figure 4D) and CD127 (IL-7R) (Figure S2B) was impaired by the absence of P2RX7; IL-7R signaling controls T cell homeostasis (Rathmell et al., 2001). Notably, most *P2rx7*^-/-^ CD4^+^ T cells showed a Th0 phenotype when differentiated in the Th1 condition.

**Figure 4.**
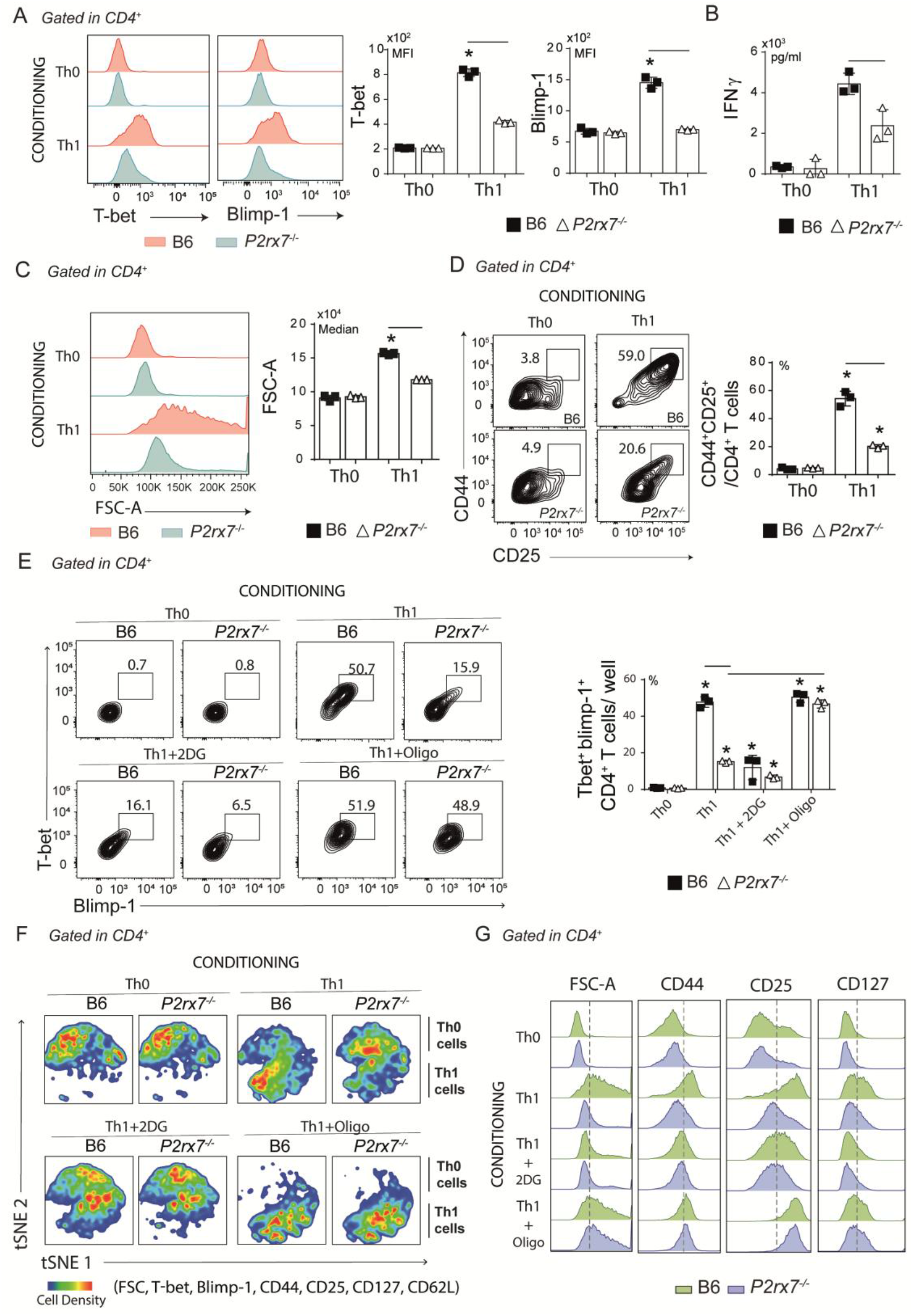
Effects of glycolytic pathway and ATP synthase inhibition *in vitro* on Th1-conditionated CD4^+^ T cells that express or not P2RX7. Splenic naïve CD4^+^ T cells from B6 and *P2rx7*^-/-^ mice were activated *in vitro* under Th1 conditions. Th0 condition was used as control. (A) Histograms show T-bet and Blimp-1 expression in CD4^+^ T cells under Th0 or Th1 conditions. Column bar graphs show T-bet and Blimp-1 MFIs. (B) Column bar graph shows IFNγ production. (C) Histogram shows FSC-A of CD4^+^ T cells and the medians are shown in the column bar graphs. (D) Contour plots show CD44 *versus* CD25 expression in CD4^+^ T cells. Column bar graph shows CD44^+^CD25^+^ T cell frequencies. (E) Contours plots show T-bet and Blimp-1 expression of in Th0 or Th1 cells in the absence or presence of 2DG or oligomycin (Oligo). Column bar graph shows T-bet^+^Blimp-1^+^CD4^+^ T cell frequencies per well. (F) The t-distributed stochastic neighbor embedding (t-SNE) maps show CD4^+^ T cell clusters from concatenated samples in relation to cell density. (G) Histograms show FSC-A and CD44, CD127 and CD25 expression in CD4^+^ T cells. Data are shown as the mean ± SD (n = 3) of one representative experiment out of three. Significant differences were observed for the (-) indicated groups with p < 0.05, using Mann Whitney U test.

The role of metabolic reprogramming in the differentiation of Th1 cells that expressed or did not express P2RX7 was then investigated by inhibiting the glycolytic pathway or oxidative phosphorylation. Glycolysis inhibition with 2-deoxi-d-glucose (2DG) reduced the expression of Blimp-1 and T-bet, most evidently for Th1-polarized B6 CD4^+^ T cells that became phenotypically similar to *P2rx7*^-/-^ CD4^+^ T cells differentiated in the same condition (Figure 4E). Remarkably, blocking ATP synthase with oligomycin that inhibits the oxidative phosphorylation and drives cellular metabolism for aerobic glycolysis was sufficient to promote Blimp-1 and T-bet expression in *P2rx7*^-/-^ CD4^+^ T cells. T-distributed stochastic neighbor embedding (t-SNE) analysis for T-bet, Bcl6, Blimp1, CD44, CD25, CD127 and FSC-A clearly demonstrated the major effect of P2RX7-mediated metabolic reprograming on Th1 differentiation (Figure 4F). Treatment with 2DG impaired the proliferative response of Th1-polarized B6 CD4^+^ T cells, as evidenced by the reduction in cell size (FSC-A) and expression of CD44, CD25 and CD127; this phenotype was indistinguisheble from that of untreated *P2rx7*^-/-^ CD4^+^ T cells differentiated in the Th1 condition (Figure 4G). Notably, ATP synthase blockade increased the proliferative response and expression of CD44, CD25 and CD127 in *P2rx7*^-/-^ CD4^+^ T cells differentiated in the Th1 condition to levels similar to those observed for Th1-polarized B6 CD4^+^ T cells.

We concluded that P2RX7-mediated metabolic reprogramming of CD4^+^ T cells for aerobic glycolysis is a general phenomenon, not restricted to malaria, which is a key event for Th1 differentiation (Figure 5). Inducing the production of ATP synthase inhibitors is likely a molecular mechanism by which P2RX7 signaling in CD4^+^ T cells upregulates aerobic glycolysis and, therefore, Th1 differentiation.

**Figure 5.**
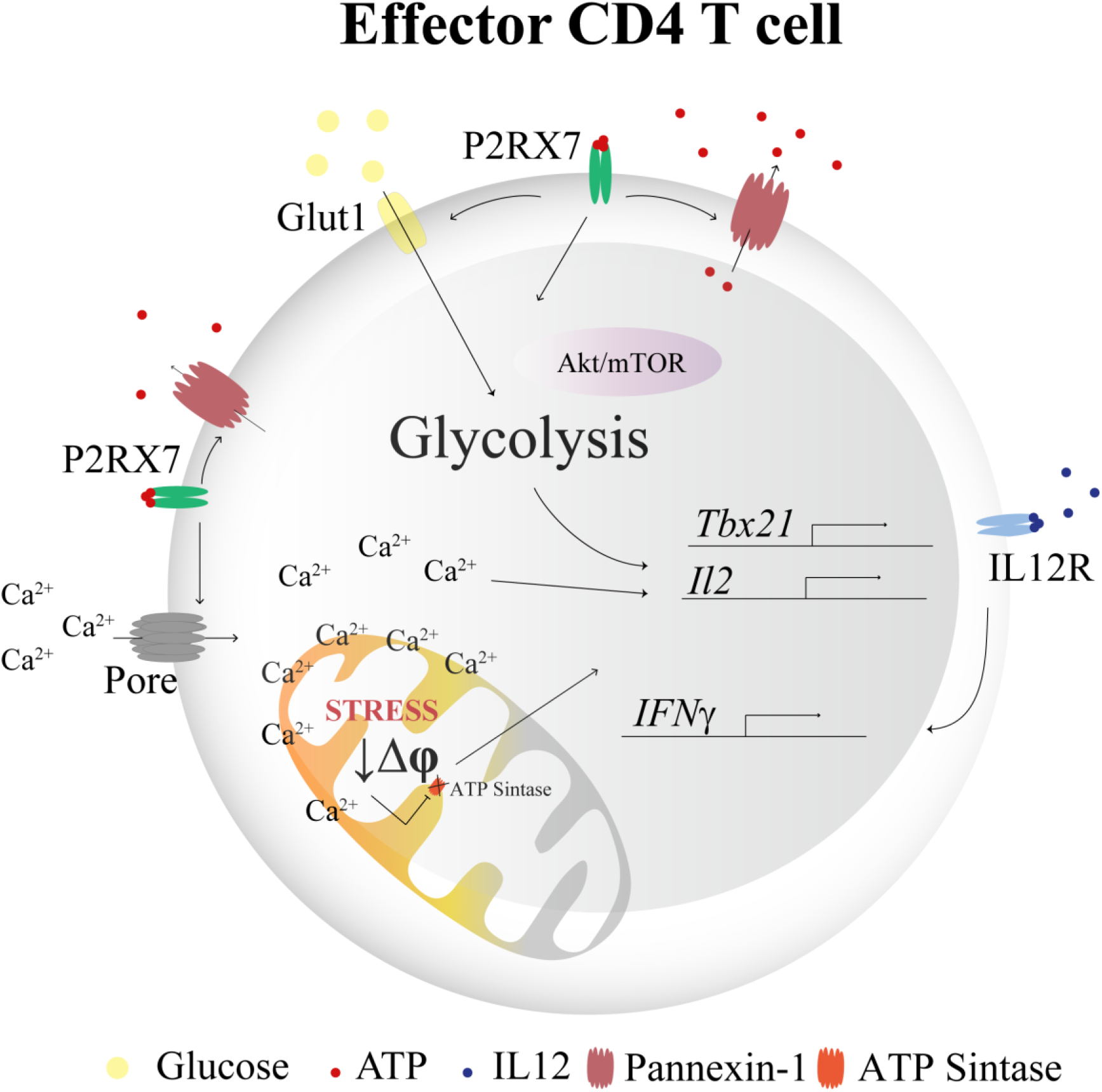
Schematic illustration showing the role of P2RX7 in Th1 differentiation through metabolic reprogramming. High extracellular ATP concentrations activate P2RX7 under stress conditions during inflammation. The intracellular ATP may be released to the extracellular space through Pannexin-1 (Panx1) channels to maintain autocrine purinergic activation. Activation of P2RX7 provides metabolic reprogramming by increasing the glucose uptake and glycolytic pathway-related proteins. P2RX7-dependent pore formation promotes high Ca^2+^ influx that leads to mitochondrial stress and mitochondrial membrane potential reduction (Δψ). In addition, Ca^2+^ influx also contribute to *il-2* transcription and proliferation. P2RX7 increase the expression of phosphorylated mTOR and Akt that culminates in t-bet transcription. ATP synthase inhibition is likely a downstream effect of P2RX7 signaling that potentiates the Th1 response. Under Th1 condition and activation of IL-12R, T-bet^+^CD4^+^ T cells produce IFNγ.

## DISCUSSION

This study describes a sequence of events arising from P2RX7 signaling in CD4^+^ T cells that respond to *Plasmodium* infection, which, together with the data published in this malaria experimental model, provides a clear view on the role of this signaling pathway in Th1 differentiation. P2RX7 acts at the beginning of the malaria immune response, before Th1/Tfh polarization, when activated CD4^+^ T cells constitute a single population that co-expresses T-bet and Bcl6, and contributes exclusively to the differentiation of Th1 cells, without affecting the generation of early Tfh cells. P2RX7 amplifies the Th1 response to malaria by inducing metabolic reprogramming for aerobic glycolysis, which supplies the anabolic intermediates needed to support rapid CD4^+^ T cell growth and proliferation (Carr et al., 2010). This was evidenced by its involvement in mTOR and Akt phosphorylation, glucose uptake and production of glycolytic enzymes such as hexokinase 1, triosephosphate isomerase 1, fructose-bisphosphatealdolase A and alpha-enolase. Supporting this, P2RX7 signaling is a key modulator of cancer cell metabolism, by increasing glucose uptake, glycolytic enzymes and glycogen stores (Amoroso et al., 2012). In contrast, Tfh cells are less proliferative and glycolytic than Th1 cells due to the lack of IL-2 signaling and mTOR and Akt activation (Ray et al., 2015) and, thus, do not require P2RX7 to differentiate.

The induction of the glycolytic pathway by P2RX7 presumably occurs during the interaction of CD4^+^ T cells with activated phagocytes, which, as previously reported, is a determinant event for Th1 differentiation in *Plasmodium* infection (Lönnberg et al., 2017). In contrast, the decision to differentiate in the Tfh phenotype relies on the interaction of activated CD4^+^ T cells with B cells in the primary lymphoid follicles (Lönnberg et al., 2017), and occurs independently of P2RX7 signaling (Salles et al., 2017). Thus, the most likely scenario is that part of the activated CD4^+^ T cells interacts with red pulp phagocytes, which are strongly stimulated by iRBCs and thus release high amounts of ATP through pannexin-1; the engagement of P2RX7 in CD4^+^ T cells induces the aerobic glycolysis and potentiates the Th1 response. Red pulp phagocytes include macrophages, monocytes and a large population of dendritic cells that actively uptake infected red blood cells (iRBCs) and interact with CD4^+^ T cells during acute malaria (Borges da Silva et al., 2015a; Borges da Silva et al., 2015b); the optimization of phagocyte functional activities by IFNγ helps to control the parasite (Bastos et al., 2002). ATP released by parasites through ion channels (Akkaya et al., 2009; Levano-Garcia, et al., 2010) or upon erythrocyte lysis may also contribute to activating P2RX7 in CD4^+^ T cells that interact with red pulp phagocytes. This view is supported by a recent study showing that P2RX7 expression in CD4^+^ T cells is particularly important for cell migration, proliferation and production of IFNγ in the pulmonary parenchyma during severe tuberculosis and influenza (Santiago-Carvalho et al., 2022).

One of the main effects of P2RX7 signaling on activated CD4^+^ T cells is to enhance the efficiency of Ca^++^ handling by the endoplasmic reticulum and, therefore, improve Ca^++^-dependent NFAT translocation to the cell nucleus, leading to IL-2 production and IL-2R expression (Adinolfi et al., 2005; Adinolfi et al., 2009). In agreement with this, P2RX7 signaling in CD4^+^ T cells increases intracellular Ca^++^ concentration and IL-2 production during *Plasmodium* infection (Salles et al., 2017), as well as induces mTOR and Akt phosphorylation as shown in this study. These metabolic sensors upregulate the expression of several transcriptional factors that are critical for the expression of glycolytic enzymes such as hypoxia-inducible factor 1α (HIF1α), interferon regulatory factor-4 (IRF-4) and c-Myc (Vaeth et al., 2017). Increased ROS production and mitochondrial depolarization are also consequences of P2RX7 activation in CD4^+^ T cells during *Plasmodium* infection, being indicative of mitochondrial Ca^++^ overload. Although mitochondrial ROS is critical for NFAT activation and IL-2 production (Kamiński et al., 2010; Sena et al., 2013), uncontrolled Ca^++^ influx from sustained P2RX7 signaling results in mitochondrial Ca^++^ overload, and therefore in exacerbated ROS production and collapse of the MMP due to opening of the mitochondrial permeability transition pore (MPTP) (Di Virgilio, et al., 2017). In this context, the selective expression of CD39 in Th1 cells (Salles et al., 2017), as well as the fact that Entpd1 (CD39 gene) is one of highest-ranking Th1 transcribed genes during Th1/Tfh subset bifurcation (Lönnberg et al., 2017), indicates that tight regulation of eATP levels is crucial to minimize the undesirable effects of excessive P2RX7 signaling in Th1 responses.

More details on the critical role of P2RX7 signaling in the metabolic reprogramming of Th1 cells are given by *in vitro* experiments in which CD4^+^ T cells are stimulated with anti-CD3 and anti-CD28 antibodies in the presence of IL-2, IL-12 and anti-IL-4 antibody. P2RX7 deficiency in CD4^+^ T cells prevents Th1 polarization, as well as pharmacological inhibition of the glycolytic pathway. Furthermore, the blockade of ATP synthase activity and consequent inhibition of oxidative phosphorylation, which drives cellular metabolism for aerobic glycolysis, is sufficient to induce IL-2R and IL-7R expression, CD4^+^ T cell proliferation and polarization to the Th1 profile in the absence of P2RX7-mediated Ca^++^ influx. These results show how glucose metabolism integrates with TCR, CD28, IL-2 and IL-12 signaling in order to ensure the differentiation of Th1 cells, in addition to highlighting the important role of P2RX7 as a catalyst for the metabolic shift to aerobic glycolysis. The fact that inhibition of ATP synthase has no important effect on polarized Th1 cells expressing P2RX7 is consistent with a previous study showing that proliferation of effector T cells does not depend on mitochondrial ATP production, although it is critical during the early activation of unpolarized T cells (Chang et al., 2013).

Our findings rise the possibility that inducting the production of ATP synthase inhibitors is a fundamental mechanism by which P2RX7 signaling promotes Th1 cell differentiation. The ATPase inhibitory factor 1 (IF1) is a master regulator of cell metabolism that once dephosphorylated inhibits oxidative phosphorylation and upregulates aerobic glycolysis (García-Bermúdez and Cuezvado, 2016). Dephosphorylated IF1 is highly expressed in colon, lung and breast carcinomas and in hypoxia (Garcia-Bermudez et al., 2015); the overexpression or silencing of IF1 results in upregulation or downregulation of glycolysis in cancer cells, respectively (Sanchez-Cenizo et al., 2010; Formentini et al., 2012; Sanchez-Arago et al., 2013). In addition, the TCA cycle metabolite α-ketoglutarate (Chin et al., 2014) and the oncometabolite (R)-2-hydroxyglutarate inhibit ATP synthase (Fu et al., 2015), as well as metabolites accumulated in some human pathologies such as neuronal ceroid lipofuscinosis and methylmalonic and glutaric acidurias (Das et al., 2003). The induction of ATP synthase inhibitors by P2RX7 may provide the optimal metabolic conditions for CD4^+^ T cell proliferation and differentiation to the Th1 profile.

This study demonstrates that P2RX7-mediated metabolic reprogramming for aerobic glycolysis is a general phenomenon, which occurs both *in vivo* during acute *P. chabaudi* malaria and in Th1-polarized *vitro* conditions, being a key event for the development of an efficient Th1 response in environments with high eATP levels. The molecular mechanism involved in this process should be better clarified from the evidence that points to the participation of ATP synthase inhibitors. Understanding the metabolic pathways involved in Th1 responses under intense stimulation can help to develop new therapeutic approaches for infectious, autoimmune and other inflammatory diseases.

## EXPERIMENTAL PROCEDURES

### Mice, parasite and infection

Six-to-eight-week-old C57BL/6 (B6), B6.129P2-*P2rx7^tm1Gab^*/J (*P2rx7*^-/-^), Foxp3.GFP, FoxP3.GFP^+^ *P2rx7 ^tm1Gab^*/J (Foxp3.GFP^+^ *P2rx7*^-/-^), B6.SJL-Ptprca Pepcb/BoyJ (CD45.1^+/+^), and *B6.129S2-Cd4tm1Mak/J* (*CD4*^-/-^) female mice (originally from The Jackson Laboratory) were bred under specific pathogen-free conditions at the Isogenic Mice Facility (ICB-USP, Brazil). The *P2rx7*^-/-^ mice were generated by Pfizer Inc. (USA). Because the schizogonic cycle of *P. chabaudi* depends on the host circadian rhythm, mice were maintained under an inverted light/dark cycle for at least 15 days before infection to access the period adjacent to erythrocyte invasion. Mice were infected intraperitoneally with 1 × 10^6^ iRBCs. All experimental procedures were in accordance with national regulations of ethical guidelines for mouse experimentation and welfare of the Health National Council and Animal Experimentation Brazilian College (COBEA) - Brazil, the protocols being approved by the Health Animal Committee of USP, with permit number 55/2017.

### Cell phenotyping

Spleen cells were washed and maintained in ice-cold RPMI 1640 supplemented with penicillin (100 U/ml), streptomycin (100 μg/ml), 2-mercaptoethanol (50 μM), L-glutamine (2 mM), sodium pyruvate (1 mM) and 3% heat-inactivated fetal bovine serum (FBS). All supplements were purchased from Life Technologies (USA). Cells (1.0 × 10^6^/well) were surface labeled with fluorochrome-conjugated monoclonal antibodies (mAbs) to CD4 (H129.19 or GK1.5), CD11a ((M17/4), CD25 (PC61), CD44 (IM7), CD127 (A7R34) (BD Bioscience, USA) and CD45.1 (104), CD45.2 (A20) (eBioscience, USA). For intracellular labeling, splenocytes (1.0 × 10^6^/well) were fixed with Cytofix/Cytoperm buffer (BD Bioscience) and permeabilized with PermWash (BD Bioscience). Cells were labeled with fluorochrome-conjugated mAbs to Blimp-1 (6D3), T-bet (eBio4B10), Bcl6 (K112-91), FoxP3 (FJK-16s), Akt (SDRNR), mTOR (MRRBY), Ki67 (B56) and IFNγ (XMG-1.2). In some experiments, cells were also stained with TMRE (100 nM, Invitrogen, USA), DCFDA (10 μM, Invitrogen), 2-NBDG (10 μM, Invitrogen) and MTG (50 nM, Invitrogen), for 15 min at 37°C, to assess MMP, ROS, glucose uptake and mitochondrial mass, respectively. Samples were acquired using FACS Fortessa or FACSCanto BD flow cytometers and the data analyzed with the FlowJo program.

### CD4^+^ T cell proliferation assay

Spleen cells (3.0 × 10^7^) were stained with 5 μM 5,6-carboxyfluorescein succinimidyl ester (CFSE; Molecular Probes, USA) in phosphate-buffered saline with 0.1% bovine serum albumin (Sigma-Aldrich, USA). CD4^+^ T cells (5 × 10^5^) were cultured with iRBCs (4 × 10^6^) for 72 h at 37°C in a 5% CO_2_ atmosphere, stained with PE-labeled mAb to CD4 and analyzed by flow cytometry.

### CD4^+^ T cell purification and adoptive transfer

CD4^+^ T cells were negatively selected using an EasySep Mouse™ CD4^+^ T Cell Isolation Kit (Stem Cell Technologies, Canada). *Cd4*^-/-^ mice were adoptively co-transferred intravenously (i.v.) with purified B6 and *P2rx7*^-/-^ CD4 T cells (1 × 10^6^/each population) and infected i.v. with 1 × 10^6^ iRBCs.

### *In vitro* Th1 differentiation and drug treatment

Splenic naïve CD4^+^ T cells were negatively selected using an EasySep™ Mouse Naïve CD4^+^ T Cell Isolation Kit (Stem Cell Technologies). Naïve T cells were resuspended in supplemented RPMI 1640 medium with penicillin (100 U/ml), streptomycin (100 μg/ml), 2-mercaptoethanol (50 μM), L-glutamine (2 mM), sodium pyruvate (1 mM), 10% heat-inactivated fetal bovine serum (FBS) and 1X MEM Non-Essential Amino Acids Solution (NEAA) (Life Technologies) and cultured in 96-well flat bottom plates at 1.0 × 10^5^/well in the presence of Dynabeads Mouse T-Activator CD3/CD28 (4×10^7^ beads/ml) (Invitrogen), 5 μg/ml of anti-IL-4 (11B11) mAb (Biolegend, USA), 5 ng/ml of IL-2 recombinant (eBioscience) and 15 ng/ml of IL-12 (eBioscience), at 37°C and 5% CO2 atmosphere for 96 h. In some cultures, cells were treated with 10mM 2-DG (Sigma-Aldrich) or 10 μM oligomycin (Sigma-Aldrich).

### Extracellular flux analysis (Seahorse)

Splenic CD4^+^ T cells were negatively selected using an EasySep Mouse™ CD4^+^ T Cell Isolation Kit (Stem Cell Technologies). CD4^+^ T cells (4.0 × 10^5^/well) were resuspended in RPMI 1640 media supplemented with 2 mM L-glutamine, 1 mM sodium pyruvate (Life Technologies) and 10 mM glucose (Sigma-Aldrich). OCR and ECAR were measured using a 96-well Seahorse XF96 bioanalyzer (Agilent, USA). In some experiments, 1 μg/ml olygomycin (Sigma-Aldrich), 5 μM FCCP (Sigma-Aldrich), 1 μg/ml antimycin A (Sigma-Aldrich), 25 mM glucose (Sigma-Aldrich) and 20 mM 2-DG (Sigma-Aldrich) were injected to obtain maximal respiratory and control values. Assay parameters were as follows: 3 min mix, no wait, 3 min measurement, repeated 3 times at basal and after each addition.

### Proteomic analysis

Splenic CD11a^+^CD4^+^ and CD11a^-^CD4^+^ T cells were separated by sorting (FACs Aria device, BD Biosciences) and frozen in a freezer −20°C (Figure S3). For protein extraction, unfreezed cells were centrifuged (700g for 3 min, 4°C) and supernatants were discarded. After washing, the pellet was resuspended in 150 μl of 8M urea buffer containing 1x protease inhibitor cocktail (Merck, Germany). It was performed 6 cycles of thermal shock: the tubes were emerged in liquid nitrogen for 30s, transferred to a 30°C bath and mixed for 1 min. Protein was quantified using Qubit (Invitrogen) and ammonium bicarbonate was added to a final concentration of 50 mM. Proteins were reduced with dithiothreitol (Sigma-Aldrich) to a final concentration of 10 mM, incubated for 45 min at 30°C, subsequently alkylated with iodoacetamide (Sigma-Aldrich) to a final concentration of 5 mM and incubated for 15 min at 30°C. Next, the samples were diluted in urea 10x by adding ammonium bicarbonate 50 mM, pH 7, to perform trypsin (Sequencin Grade Modified Trypsin) (Promega Corporation, USA) digestion at a 1:50 enzyme:protein ratio (w/w), overnight at 30°C. The samples were acidified with trifluoroacetic acid to a final concentration of 1% and digested peptides were desalted using C18 microcolumns (10.1007/978-1-61779-148-2_21). The peptides were resuspended in 0.1% Trifluoroacetic acid (TFA; protein sequencer grade), injected, and loaded on ReproSil-Pur C18 AQ (Ammerbuch-Entringen, Germany) in-house packed trap column (2 cm x 100 μm inner diameter; 5 μm). Peptides were separated at a flow of 300 nl/min on an analytical ReproSil-Pur C18 AQ packed in a house column (17 cm x 75 μm; 3 μm) by reversed-phase chromatography, which was operated on an EASY NanoLC II system (Thermo Scientific). The mobile phase was water/0.1% FA (A) and 95% ACN/0.1% FA (B). The gradient was 2-30% phase B (0.1% formic acid, 95% acetonitrile) for 80 minutes, 30-95% B for 5 minutes, and holding at 95%B for 20 minutes. The EASY NanoLC II was coupled into an LTQ Orbitrap Velos mass spectrometer (Thermo Scientific) operating in positive mode ion. The mass spectrometer acquired a full MS scan at 60,000 full width half maximum (FWHM) resolution with a 350-1500 Da mass range. The top 20 most intense peptide ions were selected from MS for Collision Ion Dissociation (CID) fragmentation (normalized collision energy: 35.0 V).

### Proteomics data processing

Raw files were processed in the Maxquant software (version 1.5.3.8) using the Andromeda search engine in the Maxquant environment. Raw data were searched against *Mus musculus* reviewed Uniprot database (November 2018 release, 17,001 entries). The parameters used were trypsin full specificity; two missed cleavages allowed, protein common contaminants, and reverse sequences included. Carbamidomethylation of cysteine as a fixed modification; methionine oxidation, and protein N-terminal acetylation as variable modifications. Identification of protein was accepted with less than 1% false discovery rate (FDR).

### Statistical analysis

Statistical analysis was performed by the Mann-Whitney U test to compare two groups. For more than two groups, data were analyzed by Kruskal-Wallis test. Survival curves were analyzed by the log-rank test using the Kaplan-Meier method. GraphPad Prism 6 software was used, in which differences between groups were considered significant when *p*< 0.05 (5%).

Proteomic statistical analysis was performed in Perseus, using Student’s T-test followed by Benjamini-Hochberg FDR correction (Tyanova et al., 2016 - https://doi.org/10.1038/nmeth.3901). Proteins identified in at least 3 samples in total and regulated with FDR<0.05 were considered for further analyses. The mass spectrometry proteomics data have been deposited to the ProteomeXchange Consortium via the PRIDE (PubMed ID: 30395289) partner repository with the dataset identifier.

## ACKNOWLEDGMENTS

We are grateful to Rogério Silva do Nascimento, Silvana Aparecida da Silva, and Maria Áurea de Alvarenga for their technical assistance, and the Central Facilities and Research Support (CEFAP) for cell sorting and proteomic analysis. This work was supported by Research Support Foundation of the State of São Paulo (FAPESP) (M.R.D.L.: 2017/03354-9 and 2015/20432-8) and National Council for Scientific and Technological Development (CNPq) (M.R.D.L.: 408909/2018-8 and 303810/2018-1).

## AUTHOR CONTRIBUTIONS

E.M.S. and M.R.D.L conceived the project and experiments. M.R.D.L. and J.M.A. provided equipments and reagents. E.M.S. and P.H.L.R. performed experiments and analyzed the data. E.M.S., C.A., V.F.S. and G.P. performed proteomic analysis. E.M.S., C.N.S., T.R. and N.O.S.C performed extracellular flux analysis (Seahorse). E.M.S. and M.R.D.L. wrote the manuscript.

## SUPPLEMENTARY INFORMATION

**Figure S1.**
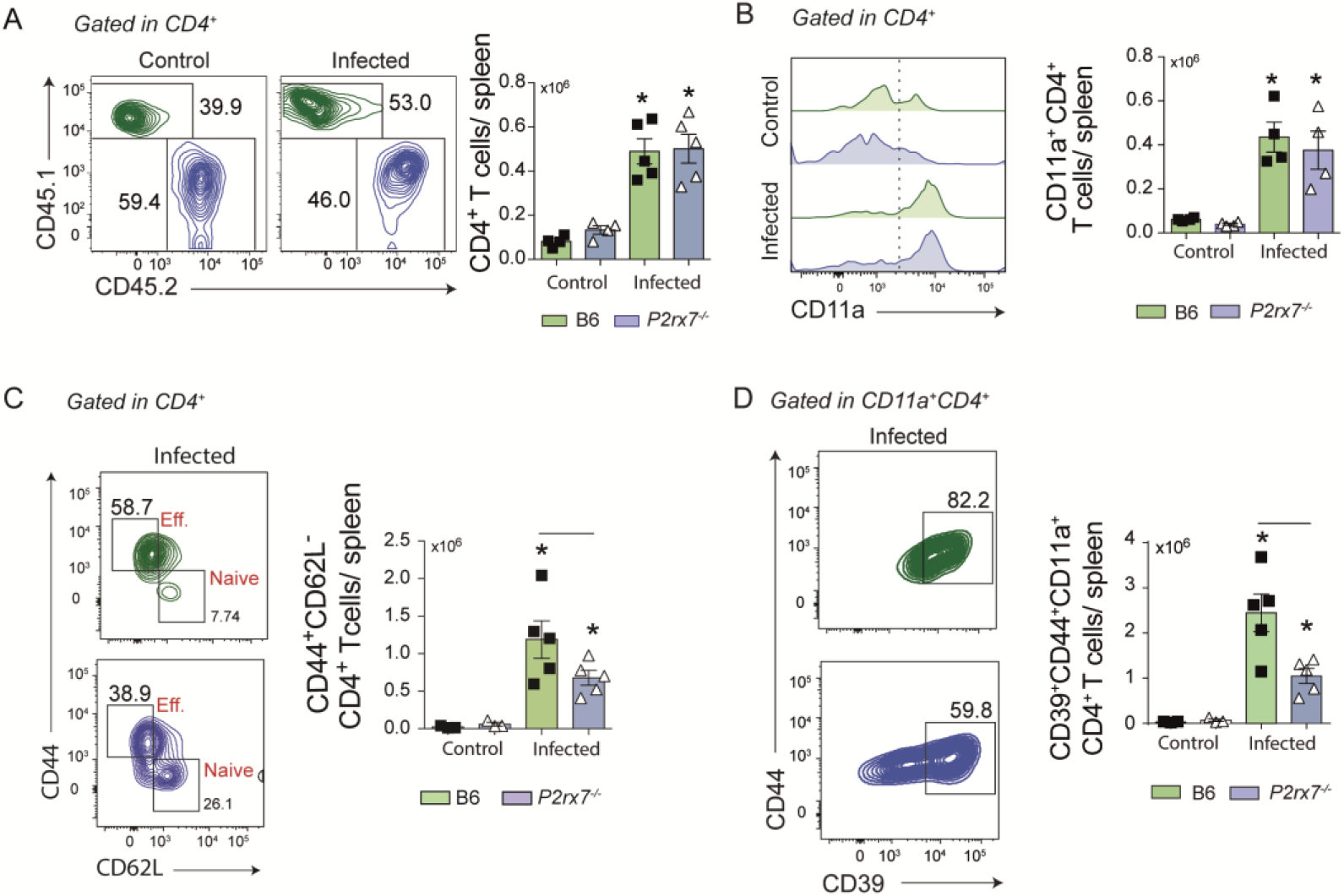
Phenotypical analysis of co-transferred B6 and *P2rx7*^-/-^ CD4^+^ T cells at early *Plasmodium* infection. Splenic naïve CD4^+^ T cells from B6 (CD45.1^+^) and *P2rx7*^-/-^ (CD45.1^+^) mice were co-transferred into *Cd4*^-/-^ mice that were infected with 1 × 10^6^ iRBCs. Splenic CD45.1^+^ and CD45.2^+^ CD4^+^ T cells were analyzed at day 6 of infection. Non-infected mice were used as controls. (A) Contour plots show CD45.1^+^ *versus* CD45.2^+^ expression in CD4^+^ T cells. Column bar graph shows CD45.1^+^ and CD45.2^+^ CD4^+^ T cell numbers per spleen. (B) Histogram shows CD11a expression in CD4^+^ T cells. Column bar graph shows CD11a^+^CD4^+^ T cell numbers per spleen. (C) Contour plots show CD44 *versus* CD62L expression in CD4^+^ T cells. Column bar graph shows CD44^+^CD62L^-^CD4^+^ (effector) and CD44^-^CD62L^+^CD4^+^ (naïve) T cell numbers per spleen. (D) Contour plots show CD44 *versus* CD39 expression in CD11a^+^CD4^+^ T cells. Column bar graph shows CD44^+^CD39^+^CD11a^+^CD4^+^ T cell numbers per spleen. Data are shown as the mean ± SD (n = 4-5) of one representative experiment out of three. Significant differences were observed for the (-) indicated groups and (*) between infected and control mice with p < 0.05, using Mann Whitney U test.

**Figure S2.**
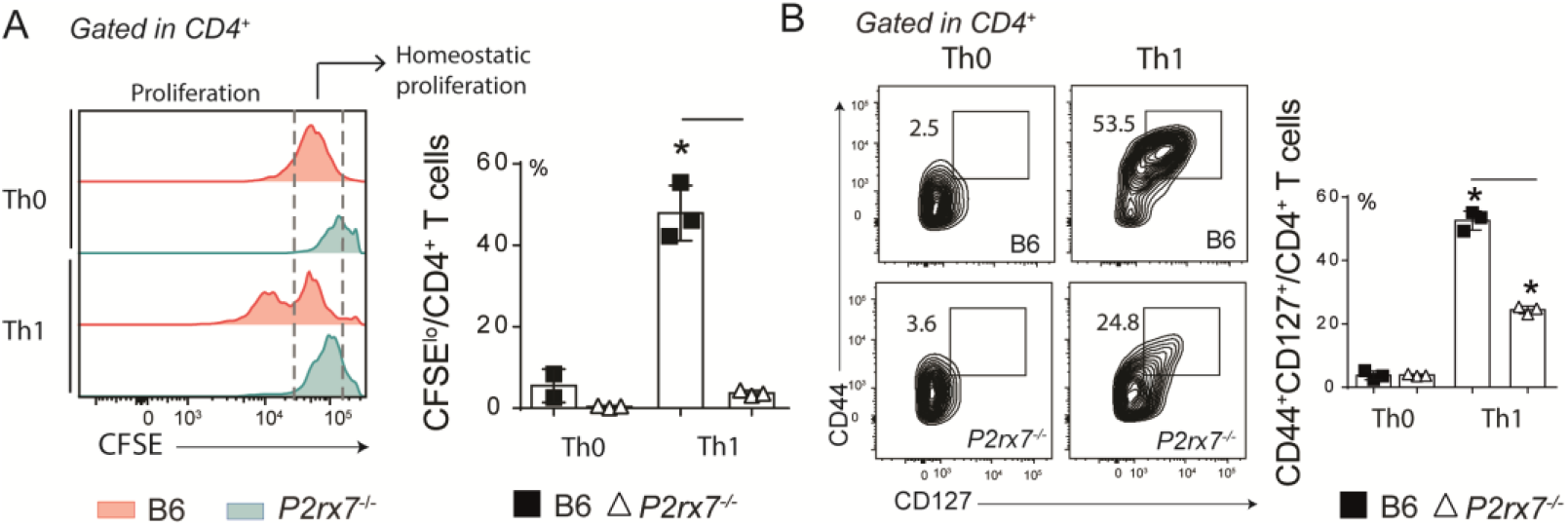
CFSE staining and CD127 expression in Th1-conditionated CD4^+^ T cells that express or not P2RX7. Splenic naïve CD4^+^ T cells from B6 and *P2rx7*^-/-^ female mice were activated under *in vitro* Th1 conditions. Th0 condition was used as control. (A) Histograms show CFSE staining CD4^+^ T cells. Column bar graph shows CSFE^lo^CD4^+^ T cell frequencies. (B) Contour plots show CD44 *versus* CD127 expression in CD4^+^ T cells. Column bar graph shows CD44^+^CD127^+^ cell frequencies. Data are shown as the mean ± SD (n = 3) of one representative experiment out of three. Significant differences were observed for the (-) indicated groups with p < 0.05, using Mann Whitney U test.

**Figure S3.**
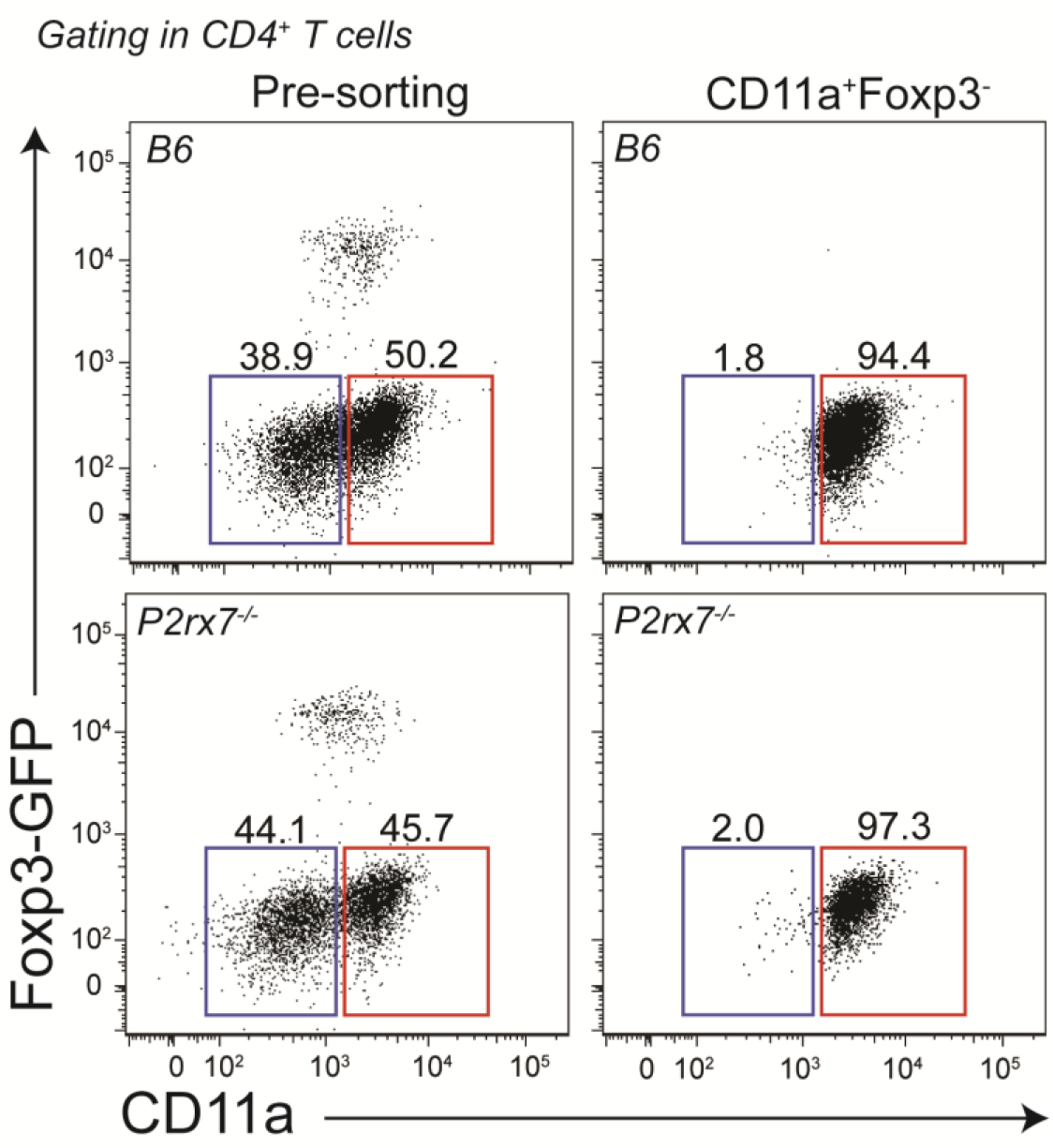
Phenotypic analysis of sorted Foxp3-CD11a+CD4+ T cells. B6 and *P2rx7*^-/-^ female mice were infected i.p. with 1 × 10^6^ iRBCs. At 6 days p.i., CD4^+^ T cells were purified using magnetic beads, and then Foxp3^-^CD11a^+^CD4^+^ T cells were sorted by BD ARIA flow cytometry. Data show one representative experiment out of six.

## REFERENCES

Adinolfi, E., Callegari, M.G., Cirillo, M., Pinton, P., Giorgi, C., Cavagna, D., Rizzuto, R., and Di Virgilio, F. (2009). Expression of the P2X7 receptor increases the Ca2+ content of the endoplasmic reticulum, activates NFATc1, and protects from apoptosis. The Journal of biological chemistry 284, 10120–10128.

Adinolfi, E., Callegari, M.G., Ferrari, D., Bolognesi, C., Minelli, M., Wieckowski, M.R., Pinton, P., Rizzuto, R., and Di Virgilio, F. (2005). Basal activation of the P2X7 ATP receptor elevates mitochondrial calcium and potential, increases cellular ATP levels, and promotes serum-independent growth. Molecular biology of the cell 16, 3260–3272.

Akkaya, C., Shumilina, E., Bobballa, D., Brand, V.B., Mahmud, H., Lang, F., Huber, S.M. (2009) The Plasmodium falciparum induced anion channel of human erythrocytes is an ATP-release pathway. Pflugers Archive 457, 1035–147.

Amoroso, F., Falzoni, S., Adinolfi, E., Ferrari, D., and Di Virgilio, F. (2012). The P2X7 receptor is a key modulator of aerobic glycolysis. Cell death & disease 3, e370.

Bastos, K.R.B., Barboza, R., Elias, R.M. Sardinha, L.R., Grisotto, M.G., Marinho, C.R.F., Amarante-Mendes, G.P., Alvarez, J.M., D’Império Lima, M.R. (2002) Impaired macrophage responses may contribute to exacerbation of blood-stage Plasmodium chabaudi chabaudi malaria in interleukin-12-deficient mice. Journal of Interferon and Cytokine Research, 22(12), 1191–1199.

Borges da Silva, H., Beura, L.K., Wang, H., Hanse, E.A., Gore, R., Scott, M.C., Walsh, D.A., Block, K.E., Fonseca, R., Yan, Y., Hippen, K.L., Blazar, B.R., Masopust, D., Kelekar, A., Vulchanova, L., Hogquist, K.A., Jameson, S.C. (2018) The purinergic receptor P2RX7 directs metabolic fitness of long-lived memory CD8^+^ T cells. Nature 559, 264–268.

Borges da Silva, H., Fonseca, R., Moreira Pereira, R., Dos Anjos Cassado, A., Álvarez, J. M., D’Império Lima, M. R. (2015) Splenic macrophage subsets and their function during blood-borne infections. Frontiers in Immunology 6, 480.

Borges da Silva, H., Fonseca, R., Dos Anjos Cassado, A., Machado de Salles, E., Nogueira de Menezes, M., Langhorne, J., Perez, K.R., Cuccovia, I.M., Ryffel, B., Barreto, V.M., Marinho, C.R.F., Boscardin, S.B., Álvarez, J.M., D’Império-Lima, M.R., Tadokoro, C.E. (2015) In vivo approaches reveal a key role for DCs in CD4^+^ T cell activation and parasite clearance during the acute phase of experimental blood-stage malaria. PLoS Pathogens, 11(2), e1004598.

Butler, N.S., Moebius, J., Pewe, L.L., Traore, B., Doumbo, O.K., Tygrett, L.T., Waldschmidt, T.J., Crompton, P.D., and Harty, J.T. (2011). Therapeutic blockade of PD-L1 and LAG-3 rapidly clears established blood-stage Plasmodium infection. Nature immunology 13, 188–195.

Cappello, P., Principe, M., Bulfamante, S., Novelli, F. Alpha-Enolase (ENO1), a potential target in novel immunotherapies. (2017) Front Biosci (Landmark Ed). 22, 944–959.

Carr, E.L., Kelman, A., Wu, G.S., Gopaul, R., Senkevitch, E., Aghvanyan, A., Turay, A.M., and Frauwirth, K.A. (2010). Glutamine uptake and metabolism are coordinately regulated by ERK/MAPK during T lymphocyte activation. Journal of immunology 185, 1037–1044.

Chang, C.H., Curtis, J.D., Maggi, L.B., Jr., Faubert, B., Villarino, A.V., O’Sullivan, D., Huang, S.C., van der Windt, G.J., Blagih, J., Qiu, J., et al. (2013). Posttranscriptional control of T cell effector function by aerobic glycolysis. Cell 153, 1239–1251.

Chi, H. (2012). Regulation and function of mTOR signalling in T cell fate decisions. Nature Reviews in Immunology 12, 325–338.

Chin, R.M., Fu, X., Pai, M.Y., Vergnes, L., Hwang, H., Deng, G., Diep, S.; Lomenick, B., Meli, V.S., Monsalve, G.C., Hu, E., Whelan, S.A., Wang, J.X., Jung, G., Solis, G.M., Fazlollahi, F., Kaweeteerawat, C., Quach, A., Nili, M., Krall, A.S., Godwin, H.A., Chang, H.R., Faull, K.F., Guo, F., Jiang, M., Trauger, S.A., Saghatelian, A., Braas, D., Christofk, H.R., Clarke, C.F., Teitell, M.A., Petrascheck, M., Reue, K., Jung, M.E., Frand, A.R., Huang, J. (2014) Themetabolite alpha-ketoglutarate extends lifespan by inhibiting ATP synthase and TOR, Nature 510 397–401.

Das, A.M. (2003) Regulation of the mitochondrial ATP-synthase in health and disease. Mol. Genet. Metab. 79 71–82.

Delgoffe, G.M., Kole, T.P., Zheng, Y., Zarek, P.E., Matthews, K.L., Xiao, B., Worley, P.F., Kozma, S.C., and Powell, J.D. (2009). The mTOR kinase differentially regulates effector and regulatory T cell lineage commitment. Immunity 30, 832–844.

Di Virgilio, F., Dal Ben, D., Sarti, A.C., Giuliani, A.L., Falzoni, S. (2017) The P2X7 Receptor in Infection and Inflammation. Immunity 47(1), 15–31.

Elias, R. M., Sardinha, L. R., Bastos, K. R. B., Zago, C. A., Freitas da Silva, A. P., Alvarez, J. M., D’Império Lima, M. R. (2005) Role of CD28 in polyclonal and specific T and B cell responses required for protection against blood stage malaria. Journal of Immunology 174(2) 790–9.

Formentini, L., Sánchez-Aragó, M., Sánchez-Cenizo, L., Cuezva, J.M. (2012) The mitochondrial ATPase inhibitory factor 1 (IF1) triggers a ROS-mediated retrograde pro-survival and proliferative response, Mol. Cell 45, 731–742.

Fu, X., Chin, R.M., Vergnes, L., Hwang, H., Deng, G., Xing, Y., Pai, M.Y., Li, S., Ta, L., Fazlollahi, F., Chen, C., Prins, R.M., Teitell, M.A., Nathanson, D.A., Lai, A., Faull, K.F., Jiang, M., Clarke, S.G., Cloughesy, T.F., Graeber, T.G., Braas, D., Christofk, H.R., Jung, M.E., Reue, K., Huang, J. (2015) 2-hydroxyglutarate inhibits ATP synthase and mTOR signaling, Cell Metab. 22 508–515.

García-Bermúdez, J., Cuezva, J.M. The ATPase Inhibitory Factor 1 (IF1): A master regulator of energy metabolism and of cell survival. (2016) Biochimica et Biophysica Acta 1857, 1167–1182.

Garcia-Bermudez, J., Sanchez-Arago, M., Soldevilla, B., Del Arco, A., Nuevo-Tapioles, C., Cuezva, J.M., PKA phosphorylates the ATPase inhibitory factor 1 and inactivates its capacity to bind and inhibit the mitochondrial H-ATP synthase, Cell Rep. 12 (2015) 2143–2155.

Gong, D., and Malek, T.R. (2007). Cytokine-dependent Blimp-1 expression in activated T cells inhibits IL-2 production. Journal of immunology 178, 242–252.

Kaminski MM, Sauer SW, Klemke CD, Süss D, Okun JG, Krammer PH, Gülow K (2010) Mitochondrial reactive oxygen species control T cell activation by regulating IL-2 and IL-4 expression: mechanism of ciprofloxacin-mediated immunosuppression. Journal of Immunology 184, 4827–41.

Levano-Garcia, J., Dluzewski, A.R., Markus, R.P., Garcia, C.R. (2010) Purinergic signalling is involved in the malaria parasite Plasmodium falciparum invasion to red blood cells. Purinergic Signal 6, 365–72.

Liao, W., Lin, J.X., Wang, L., Li, P., and Leonard, W.J. (2011). Modulation of cytokine receptors by IL-2 broadly regulates differentiation into helper T cell lineages. Nature immunology 12, 551–559.

Lönnberg, T., Svensson, V., James, K.R., Fernandez-Ruiz, D., Sebina, I., Montandon, R., Soon, M.S., Fogg, L.G., Nair, A.S., Liligeto, U., Stubbington, M.J., Ly, L.H., Bagger, F.O., Zwiessele, M., Lawrence, N.D., Souza-Fonseca-Guimaraes, F., Bunn, P.T., Engwerda, C.R., Heath, W.R., Billker, O., Stegle, O., Haque, A., Teichmann, S.A. (2017). Single-cell RNA-seq and computational analysis using temporal mixture modelling resolves Th1/Tfh fate bifurcation in malaria. Science Immunology 2, eaal2192.

MacIver, N.J., Michalek, R.D., and Rathmell, J.C. (2013). Metabolic regulation of T lymphocytes. Annual Review of Immunology 31, 259–283.

Perez-Mazliah, D., Langhorne, J. (2015) CD4 T-cell subsets in malaria: TH1/TH2 revisited. Frontiers in Immunology 5, 671.

Muxel, S.M., Freitas do Rosário, A.P., Zago, C.A., Castillo-Méndez, S.I., Sardinha, L.R., Rodriguez-Málaga, S.M., Câmara, N.O., Álvarez, J.M., Lima, M.R. (2011) The spleen CD4^+^ T cell response to blood-stage Plasmodium chabaudi malaria develops in two phases characterized by different properties PLoS One. 6, e22434.

Rathmell, J.C., Farkash, E.A., Gao, W., and Thompson, C.B. (2001). IL-7 enhances the survival and maintains the size of naive T cells. Journal of immunology 167, 6869–6876.

Ray, J.P., Staron, M.M., Shyer, J.A., Ho, P.C., Marshall, H.D., Gray, S.M., Laidlaw, B.J., Araki, K., Ahmed, R., Kaech, S.M., et al. (2015). The Interleukin-2-mTORc1 Kinase Axis Defines the Signaling, Differentiation, and Metabolism of T Helper 1 and Follicular B Helper T Cells. Immunity 43, 690–702.

Salles, E.M., Menezes, M.N., Siqueira, R., Borges da Silva, H., Amaral, E.P., Castillo-Mendez, S.I., Cunha, I., Cassado, A.D.A., Vieira, F.S., Olivieri, D.N., et al. (2017). P2X7 receptor drives Th1 cell differentiation and controls the follicular helper T cell population to protect against Plasmodium chabaudi malaria. PLoS Pathogens 13, e1006595.

Sanchez-Arago, M., Formentini, L., Martinez-Reyes, I., Garcia-Bermudez, J., Santacatterina, F., Sanchez-Cenizo, L., Willers, I.M., Aldea, M., Najera, L., Juarranz, A., Lopez, E.C., Clofent, J., Navarro, C., Espinosa, E., Cuezva, J.M. (2013) Expression, regulation and clinical relevance of the ATPase inhibitory factor 1 in human cancers, Oncogenesis 2, e46.

Sanchez-Cenizo, L., Formentini, L., Aldea, M., Ortega, A.D., Garcia-Huerta, P., Sanchez-Arago, M., Cuezva, J.M. (2010) Up-regulation of the ATPase inhibitory factor 1 (IF1) of the mitochondrial H+-ATP synthase in human tumors mediates the metabolic shift of cancer cells to a Warburg phenotype. J. Biol. Chem. 285 25308–25313.

Santiago-Carvalho, Gislane Almeida-Santos, Bruna Gois Macedo, Caio Cesar Barbosa-Bomfim, Fabricio Moreira Almeida, Marcos Vinícios Pinheiro Cione, Trupti Vardam-Kaur, Mia Masuda, Sarah Van Dijk, Bruno Marcel Melo, Rogério Silva do Nascimento, Rebeka da Conceição Souza, Alba Lucínia Peixoto-Rangel, Robson Coutinho-Silva, Mario H. Hirata, José Carlos Alves-Filho, José Maria Álvarez, Elena Lassounskaia, Henrique Borges da Silva, Maria Regina D’Império-Lima (2022) T cell-specific P2RX7 favors lung parenchymal CD4+ T cell accumulation in response to severe lung infections. bioRxiv 2022.09.19.508603.

Schenk, U., Westendorf, A.M., Radaelli, E., Casati, A., Ferro, M., Fumagalli, M., Verderio, C., Buer, J., Scanziani, E., and Grassi, F. (2008). Purinergic control of T cell activation by ATP released through pannexin-1 hemichannels. Science signaling 1, ra6.

Sena, L.A., Li, S., Jairaman, A., Prakriya, M., Ezponda, T., Hildeman, D.A., Wang, C.R., Schumacker, P.T., Licht, J.D., Perlman, H., et al. (2013). Mitochondria are required for antigen-specific T cell activation through reactive oxygen species signaling. Immunity 38, 225–236.

Siska, P.J., Kim, B., Ji, X., Hoeksema, M.D., Massion, P.P., Beckermann, K., Wu, J., Chi, J.-T., Hong, J., Rathmellet, J. (2016) Fluorescence-based measurement of cystine uptake through xCT shows requirement for ROS detoxification in activated lymphocytes. Journal of Immunological Methods 438, 51–58.

Soon, M.S.F., Lee, H.J., Engel, J.A., Straube, J., Thomas, B.S., Pernold, C.P.S., Clarke, L.S., Laohamonthonkul, P., Haldar, R.N., Williams, C.G., Lansink, L.I.M., Moreira, M.L., Bramhall, M., Koufariotis, L.T., Wood, S., Chen, X., James, K.R, Lönnberg, T., Lane, S.W., Belz, G.T., Engwerda, C.R., Khoury, D.S., Davenport, M.P., Svensson, V., Teichmann, S.A., Haque, A. (2020) Transcriptome dynamics of CD4+ T cells during malaria maps gradual transit from effector to memory. Nature Immunology 21(12), 1597–1610.

Steinberg, T.H., and Silverstein, S.C. (1987). Extracellular ATP4-promotes cation fluxes in the J774 mouse macrophage cell line. The Journal of biological chemistry 262, 3118–3122.

Steinberg, T.H., Buisman, H.P., Greenberg, S., Di Virgilio, F., and Silverstein, S.C. (1990). Effects of extracellular ATP on mononuclear phagocytes. Annals of the New York Academy of Sciences 603, 120–129.

Takenaka, M. C., Robson, S., Quintana, F. J. (2016) Regulation of the T cell response by CD39. Trends Immunol. 37, 427–39.

Vaeth, M., Yang, J., Yamashita, M., Zee, I., Eckstein, M., Knosp, C., Kaufmann, U., Karoly Jani, P., Lacruz, R.S., Flockerzi, V., et al. (2017). ORAI2 modulates store-operated calcium entry and T cell-mediated immunity. Nature communications 8, 14714.

Van der Windt, G.J., and Pearce, E.L. (2012). Metabolic switching and fuel choice during T-cell differentiation and memory development. Immunological Reviews 249, 27–42.

Woehrle, T., Yip, L., Manohar, M., Sumi, Y., Yao, Y., Chen, Y., and Junger, W.G. (2010). Hypertonic stress regulates T cell function via pannexin-1 hemichannels and P2X receptors. Journal of Leukocyte Biology 88, 1181–1189.

